# Identification and cryoEM Structure Determination of Escherichia Phage YDC107 Tail Found in a Bacteria-Contaminated Buffer

**DOI:** 10.1101/2024.12.10.627647

**Authors:** Matthew C. Jenkins, Tahiti Dutta, Daija Bobe, Mykhailo Kopylov

**Affiliations:** School of Chemistry and Biochemistry, Georgia Institute of Technology, GA, 30332, USA; New York Structural Biology Center, NY, 10044, USA

## Abstract

Cryo-electron microscopy data analysis can yield multiple structures from a single heterogeneous dataset. Here, we show a workflow we used for the identification of a contaminant from a cryoEM grid without prior knowledge of protein sequence. We determined the tail structure of Escherichia phage YDC107 from only several thousand particles. The workflow combines high-resolution single-particle data processing with *de novo* model determination using ML-based methods. Structural analysis revealed that the central part of the phage tail has a C6 symmetry, however the overall symmetry of each segment is C3 due to dimerization of a flexible domain.

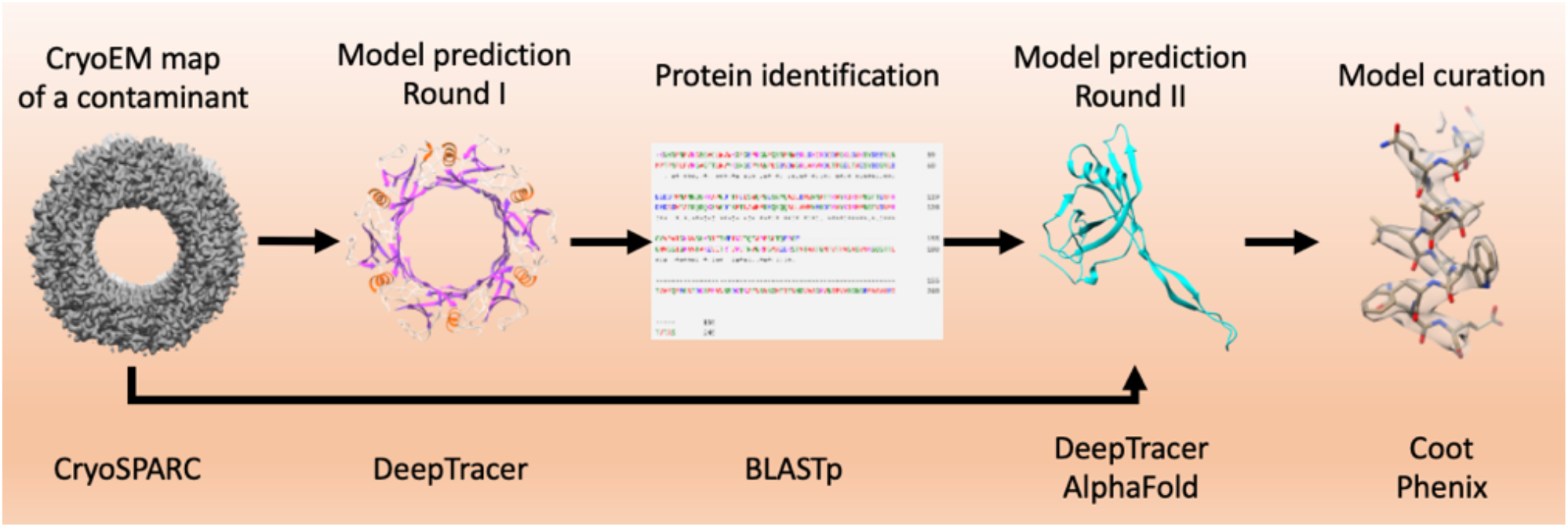

## Introduction

Cryo-electron microscopy (cryoEM) is a powerful structural biology technique due to its ability to achieve near-atomic resolution, sometimes with only several hours of data acquisition [1]. CryoEM is applicable to a wide variety of biological samples, including proteins, nucleic acids, and large macromolecular complexes like ribosomes and viruses, that can be observed in a near-native frozen-hydrated state [2,3,4].

Modern single-particle analysis (SPA) tools enable processing of highly heterogeneous samples, permitting the determination of multiple structures from a single dataset [5,6]. If sufficiently high-resolution reconstructions can be obtained, the resulting maps can be used to predict the sequence of resolved protein either manually or using such tools as Phenix map-to-model [7], DeepTracer [8] or Modelangelo [9]. Viruses, in particular bacteriophages, can be easy targets for such analyses due to the high symmetry of their building blocks such as icosahedral capsids and helically symmetric tails.

Here we used *de novo* model building to identify a protein contaminant within a recombinantly-expressed protein sample as the helical tail of the YDC107 bacteriophage. We used a combination of DeepTracer and AlphaFold [10] implementation ColabFold [11] to build the model into a 3.2 Å map. AlphaFold multimer modelling and analysis of low pass filtered maps allowed us to identify a symmetry mismatch between the hexameric core of the phage tail and the overall trimeric structure of the individual tail segments.

## Results

The original intention of the data acquisition was structure determination of a PP7 virus-like particle construct (Figure 1A, green circles) [12]. However, during grid screening, we noticed that the grid was contaminated with what appeared to be bacteria (Figure S1). Bacteria were most likely present in a compromised storage or dilution buffer. On high magnification images, we found more evidence of bacteria – bacterial flagella (Figure 1A, yellow thin arrows). We think the cells on the grids are *E*. *coli* based on their rod-like shape and size. Interestingly, we also noticed “train track” particles in many micrographs – an easily recognizable feature of bacteriophage tails (Figure 1A, blue thick arrows). We estimated that about one out of three micrographs contained the putative phage tails.

**Figure 1.**
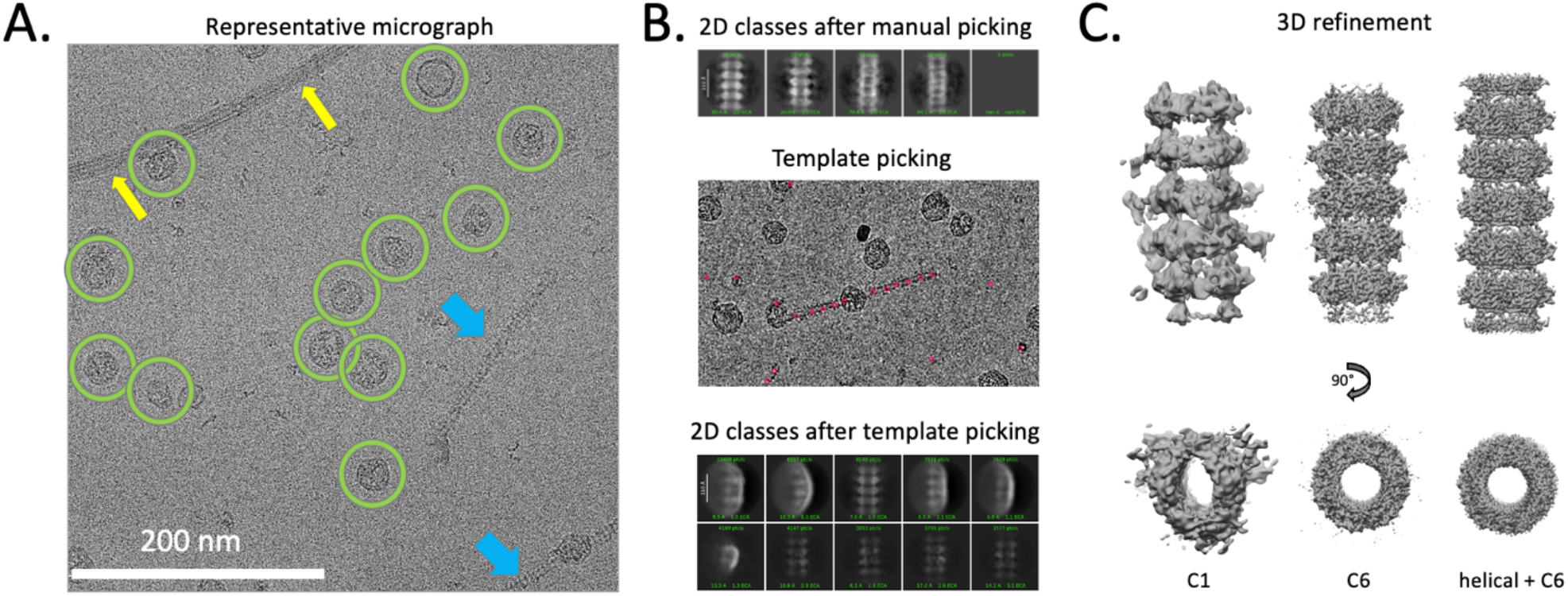
CryoEM analysis of the contaminated sample. A. CryoEM micrograph showing sample details: yellow thin arrows – bacterial flagellum; green rings – PP7 cages; blue thick arrows – fragments of bacteriophage tails. B. Particle picking and 2D classification shows an abundance of similar-looking segmented particles C. Helical 3D refinement with C6 symmetry results in 3.2 Å structure. Helical parameters were determined from a refinement with C6 symmetry.

We manually picked 154 putative bacteriophage tail segments and 2D classified them into 5 classes (Figure 1B, top). Despite such a small dataset, clear 2D averages were generated. Since each tail fragment contains multiple copies of the progenitor structural proteins, we decided to proceed with full dataset analysis. Initial 2D classes were used for template picking of the whole dataset (Figure 1B, middle), which resulted in 149,627 particles (Figure 1B, S2A). After two rounds of 2D classification, where only classes with sharp details were selected (Figure S2A), a total of 5,927 particles were used for *ab-initio* map generation and refinement (Figure 1C, Figure S2C). The resulting *ab-initio* map was used as an initial volume for C1 and C6 refinement (Figure S2 D, E). Refinement with C6 symmetry produced a 3.5 Å map and was used to estimate helical parameters. The same map was used as an initial volume for the final helical refinement with C6 symmetry (Figure 1C, Figure S3). The final map had a reported GSFSC resolution of 3.22 Å and was b-factor corrected based on the estimation from a Guinier plot (Figure S3C). This final map was used for structure prediction and model building of the phage tail’s core.

We used the online DeepTracer server to generate an initial model prediction (Figure 2A). From the predicted model, we selected one of the monomers of the central 6-member ring and extracted sequence information from it. The extracted putative sequence was used to search the NCBI database using BLASTp with default seangs without any restrictions. All top 100 hits pointed to the *Escherichia coli* phage tail protein, specifically phage YDC107 – an unclassified *Ravinvirus* from a *Caudioviricetes* class. This further supported our guess that the bacterial contamination was *E. coli*.

**Figure 2.**
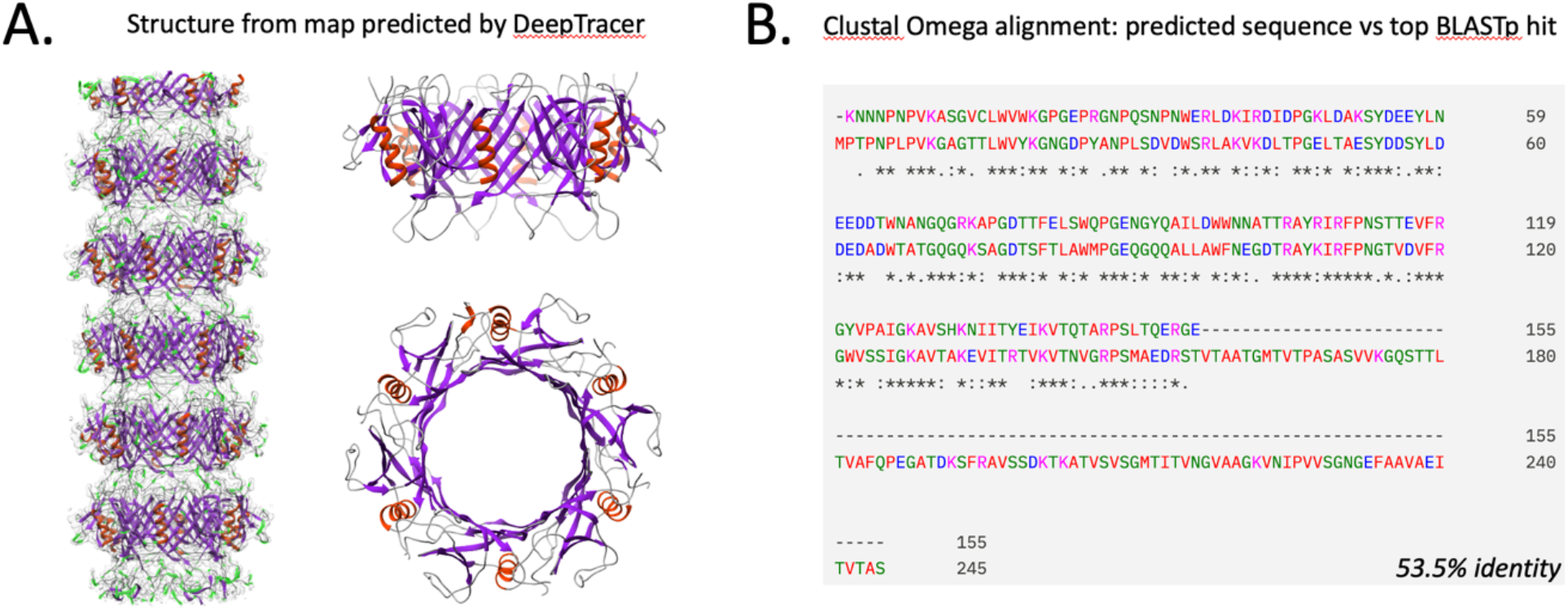
Sequence identification from cryoEM map. A. DeepTracer-predicted structure from the helically symmetrized 3.2 Å sharpened map. B. Multiple sequence alignment of the predicted sequence and the top hit from a BLASTp search, using the predicted sequence as a query.

We chose the top max score sequence with accession code MCN3709170.1 as our reference sequence. Clustal omega [13] alignment showed that about 100 amino acids on the C-terminus side were missing, while the rest shared 53.5% sequence identity (Figure 2B). This sequence was used for a second run of DeepTracer, using the same helical map but now providing it with the reference sequence identified in the BLASTp search. The output of the second run of DeepTracer was inspected in Coot [13], sequence mismatches were scrutinized, and, in all cases, reverted to match the reference sequence (Figure S4). The same reference sequence was also used to generate models using AlphaFold.

Comparison between the DeepTracer model and AlphaFold models revealed that only residues 1-154 were predicted by DeepTracer (Figure 3A). This corresponds to visual inspection of the C6-helical refined map in Coot – there were no unmodelled densities. Overall, AlphaFold and DeepTracer-derived models for the residue range [1-154] agree very well, with a RMSD between 1185 atom pairs of 3.3 Å. However, the AlphaFold model suggested the presence of an additional β-sheet domain connected to the central core by a flexible linker (Figure 3A, magenta).

**Figure 3.**
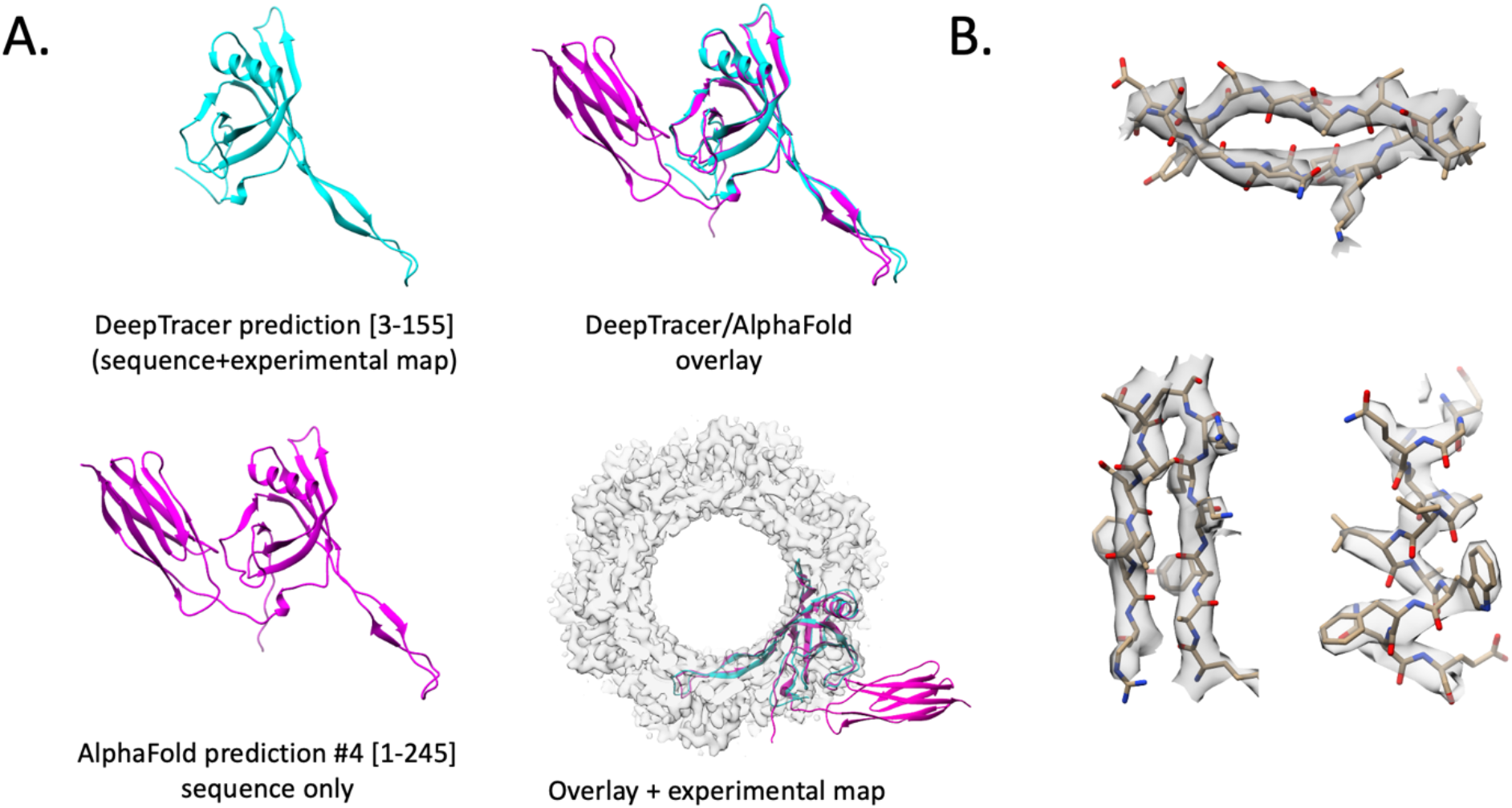
Bacteriophage YDC107 sequence validation by structure building. A. The sequence identified from the BLASTp search was used as an input to DeepTracer, and the resulting model was then manually edited in COOT residues. The same sequence was also used as an input for AlphaFold. Modelled residues with DeepTracer are 3-155 out of 245, numbering based on the reference sequence from BLASTp search: GenBank: MCN3709170.1. B. Selected map regions showing map-to-model correspondence. Antiparallel beta-sheet (top) – [106-112, 116-121], alpha-helix [91-102] (bottom right); beta-hairpin (bottom left) – [50-56, 70-77].

Low-pass filtering the C6-helical map to 12 Å resolution revealed the presence of extra protruding domains that could potentially accommodate the unmodelled flexible domain that was observed in the AlphaFold prediction (Figure 4A). Rigid-body fiang the domain into one of the protruding densities showed that there was enough room for two flexible domains. We used AlphaFold multimer to generate five models of the flexible domain dimer (Figure 4B), all of which formed a C2-symmetric arrangement and fit well into the observed unmodelled density. We chose the AlphaFold model in which the N-terminal Ser155 was positioned the closest to the C-terminal Arg154 of the central core model. Since the flexible domains formed a dimer, two additional flexible domain dimers were positioned around the central core forming an overall C3-symmetric arrangement for the tail segment (Figure 4C). Adjacent C and N termini were connected to produce a composite C3-symmetric model. The composite model was then used to generate single- and double-segment masks that were applied in 3D classification and refinement, confirming a C3 arrangement of individual tail segments (Figure 4D).

**Figure 4.**
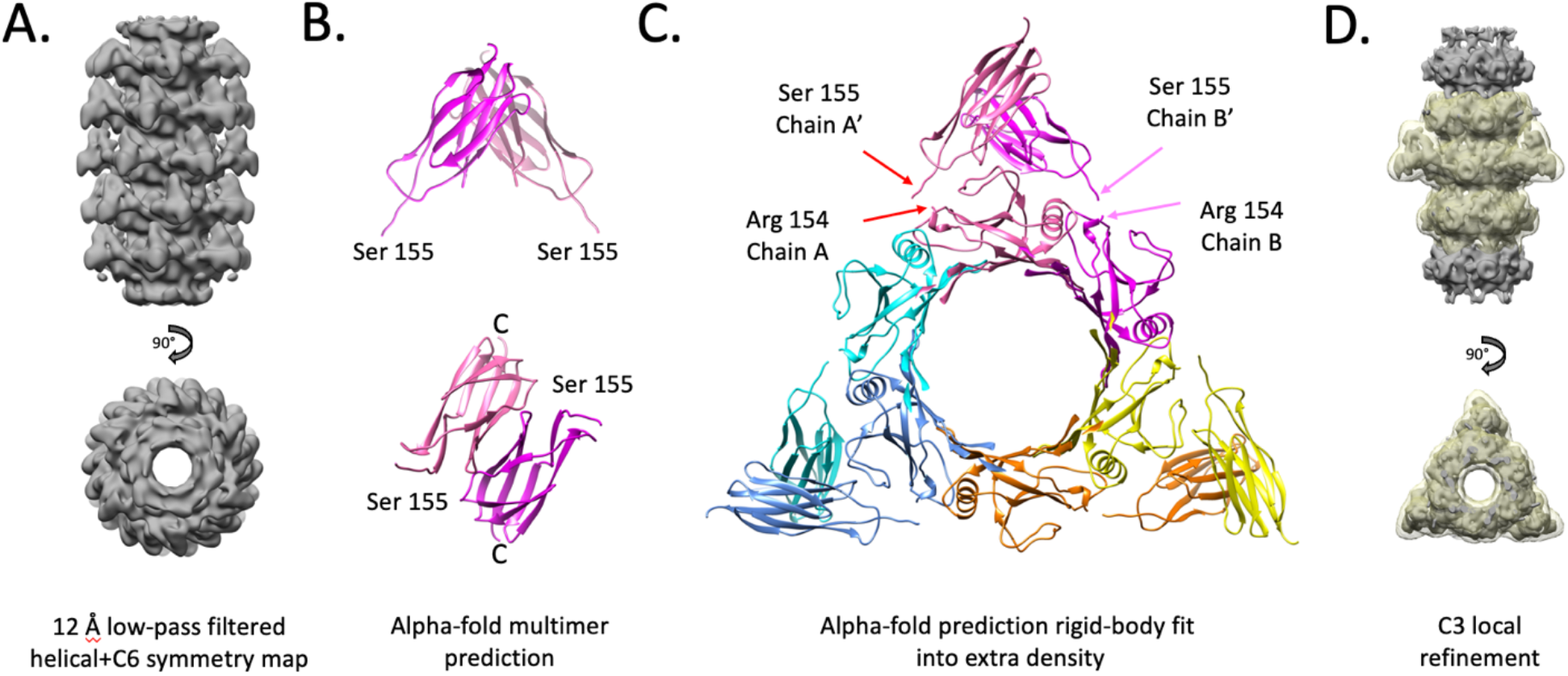
Modelling the missing domains using AlphaFold multimer. A. Helical C6 map low-pass filtered to 12 Å resolution and displayed at high threshold to show extra densities. B. Output of the AlphaFold multimer prediction model where residues 155-245 from the reference sequence were used as an input. Two domains form a C2-symmetric dimer. C. Flexible domain dimer rigid-body fitted into the extra densities of the 12 Å low-pass filtered map from panel A. The N-termini of flexible domains are positioned exactly next to the C-termini of the hexameric core model. D. Asymmetric masked 3D classification shows that there are only three extra densities per tail segment, each accommodating a flexible protein dimer.

Due to the flexible domain dimerization, individual phage tail monomers can exist in two conformations: forward and reverse (Figure 5A). The forward conformation is similar to the one predicted by AlphaFold for a full-length protein. In this conformation, the flexible domain is “reaching” forward following the direction of helical twist. In the reverse conformation, the direction of the flexible domain is opposite of the helical twist, allowing it to form a dimer with the preceding phage tail monomer that is in a forward conformation.

**Figure 5.**
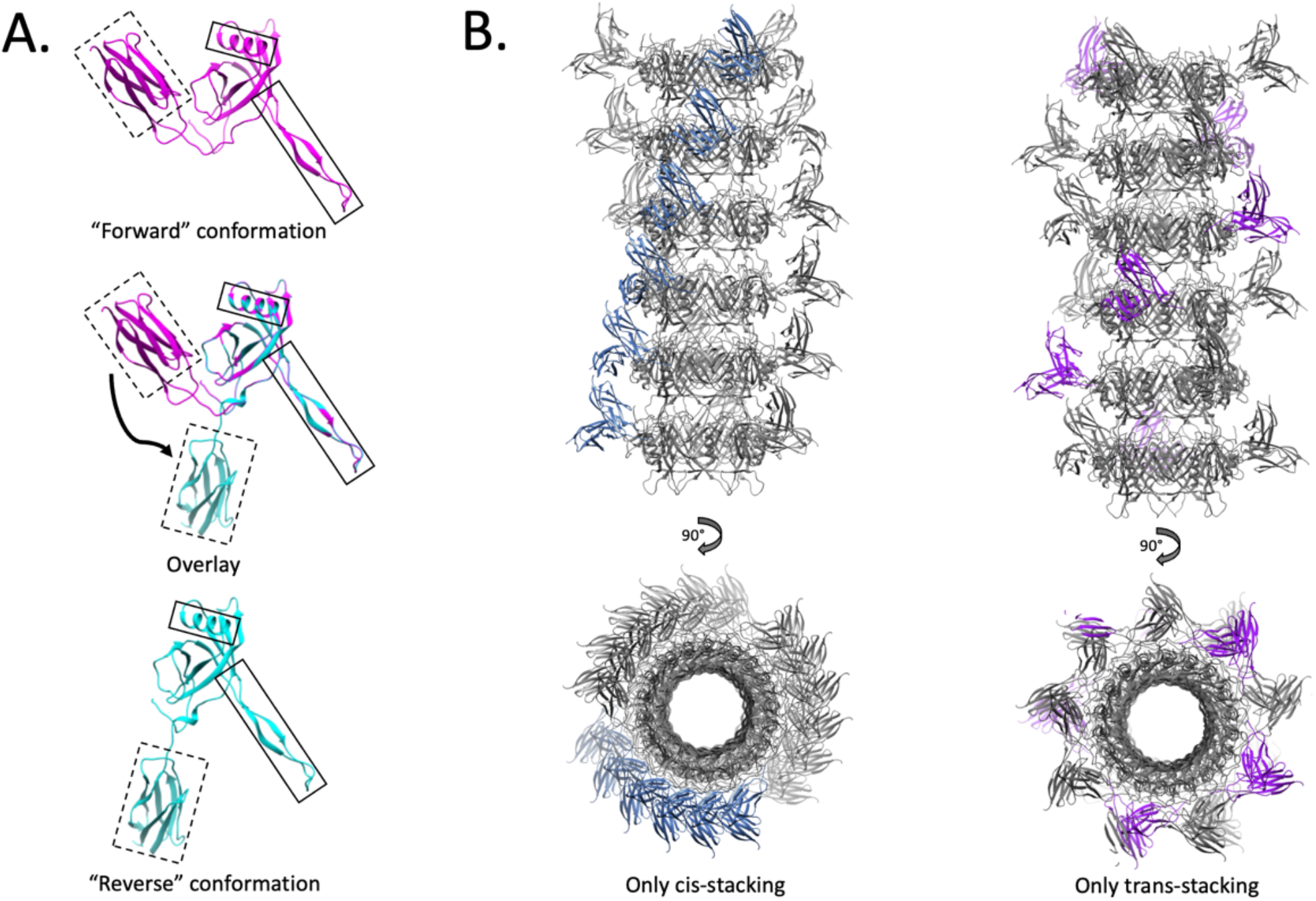
Two conformations of the phage tail protein monomer. A. Forward conformation - this model is like the one predicted by AlphaFold for the full-length protein. Reverse conformation – in this conformation the protein will interact with the flexible domain of the adjacent subunit that should be in “forward” conformation for the dimerization to occur. In the middle - overlay of forward and reverse conformations, where residues 1-155 for both models are superimposed. B.Two possible arrangements between adjacent monomers give rise to either cis-or trans-conformations. Depicted here are two tail models where all segments are either in cis-conformation (lea) or in trans-conformation (right).

Further, we hypothesized that adjacent tail segments could exist in two different conformations. To test this, we generated two kinds of masks that covered the two adjacent segments – one where flexible domain pairs are close to each other between segments, and another one where flexible domain pairs are further apart. 3D classification using these masks showed two types of stacking for adjacent tail segments: cis-stacking (Supplementary video 1) and trans-stacking (Supplementary video 2). Cis-stacked segments are offset by ∼20 degrees, which corresponds to the helical offset of the central core (Figure S5A). Trans-stacked segments are offset by ∼80 degrees, which corresponds to the helical offset of the central core plus a register shift by 60 degrees due to the 6-fold central core symmetry (Figure 5B and Figure S5B).

## Discussion

One of the initial motivations behind this study was to highlight the power of cryoEM in resolving structures from very limited datasets, without the need for prior information. In this study, we used only 5,927 segments of a helical bacteriophage tail assembly to generate a 3.3 Å map, which was sufficient for direct sequence identification from the cryoEM map. Several key features of this dataset and sample made this possible. First, due to the high symmetry, each phage tail fragment contains multiple copies of the same protein, significantly boosting signal-to-noise. In our case (6-fold cyclical symmetry), the actual number of particles used in averaging is 5,927 × 6 = ∼36,000. Second, there is very little conformational or compositional heterogeneity, at least for the central core region. Third, due to C6 and helical symmetry, this small dataset contained all the necessary views for high-resolution reconstruction, despite the absence of “top views.” Finally, each micrograph contains other particles (PP7 VLPs) that are not useful for the reconstruction of the phage tail but are critical for obtaining accurate per-micrograph estimations of the CTF.

We believe that “sequencing” from cryoEM maps for bacteriophage profiling could be a valuable method in addition to conventional nucleic acid sequencing. We recently applied this method to characterize the novel Acinetobacter baumannii phage “Mystique” [15]. In the case of Mystique, a bacterial culture lysed by the phage was applied directly to the cryoEM grid. The proteins for the tail and capsid were identified by searching the assembled viral genome using sequences obtained from the neural network prediction by Modelangelo [9] sonware.

## Materials and Methods

### Protein Expression, Purification, and Possible Sources for the Phage Contamination

Due to the source of the contamination identified on cryoEM grids is unknown, we provide the complete protocol that was used for purification of our target particles – PP7 VLPs in supplementary methods.

While the exact source of the bacterial contamination in the VLP solutions used for cryoEM cannot be explicitly identified, we speculate that it may have occurred at one of several steps in the purification pipeline. Buffers used during the purification procedure were all freshly made and sterile filtered through 0.2 μm PES filters to limit bacterial contamination. However, many of the liquid containers (e.g., beakers, graduated cylinders, conical flasks, etc.) used for storage and dilution of buffer solutions throughout the purification process are communal use items among multiple lab members that may not have been properly or effectively cleaned prior to their utilization. Additionally, we typically pass the final purified VLP solution through 0.2 μm PES syringe filters prior to usage in experimental tests. This final filtration step was inadvertently omitted for these VLP samples, and thus is the most likely source of the observed bacterial contamination.

### CryoEM Sample Preparation and Data Acquisition

Quantifoil 1.2/1.3 300 Cu grids were plasma cleaned with a hydrogen and oxygen mix for 15 seconds. A Vitrobot Mark IV was used to plunge freeze grids at 100% humidity and 22 °C by applying 3 μL of sample and using a blot force of 1 and a blot time of 6.5 s.___ Data were acquired using Leginon [14] on a NYSBC Krios4 equipped with a Gatan K3 camera and BioQuantum energy filter with a 20 eV energy slit (Table 1). A total of 3,149 movies were acquired, motion corrected, and dose-weighted using MotionCor2. Resulting images were imported to CryoSPARC for processing.

**Table 1.** Data collection and processing.

|  |  |
| --- | --- |
| Magnification | 81,000 |
| Voltage (kV) | 300 |
| Electron exposure (e/Å<sup>2</sup>) | 54 |
| Defocus range (μm) | 0.7-2.5 |
| Pixel size (Å) | 1.058 |
| Symmetry imposed | C6, helical (twist 17.90°, rise 41.77 Å) |
| No. initial particle images | 149,627 |
| No. final particle images | 5,927 |
| Map resolution FSC threshold (Å) | 3.22 |
| Map resolution range (Å) | 3.43-3.09 |

**Table 2.** Refinement.

|  |  |
| --- | --- |
| Initial model used | Ab-initio |
| Model resolution FSC threshold (Å) | 3.3 |
| Model resolution range (Å) | 3.3 |
| Map sharpening B factor (Å<sup>2</sup>) | -85.4 |
| R.m.s. deviations from ideality | From PDB validation report |
| Bonds RMSZ | 0.27 |
| Angles RMSZ | 0.53 |
| MolProbity score | From PDB validation report |
| Clashscore (all-atom clashscore) | 6 |
| Poor rotamers (%) | 4% |
| Ramachandran plot | From Phenix refinement |
| Favored (%) | 96 |
| Allowed (%) | 4 |
| Disallowed (%) | 0 |

### Data Processing

All data processing was done in CryoSPARC v.4.6.0. CTF was estimated using Patch-CTF job with default settings. 154 particles were manually picked and extracted with a box size of 256. The resulting particle stack was classified in 2D into 5 classes and the sharpest class was selected for template picking. For the template picking job, the following parameters were set: particle diameter – 192 Å, maximum number of local maxima to consider – 100, min. separation distance (diameters) – 0.5. Picked particles were extracted with a box size of 256 and cleaned up in 2D to produce a final stack of 5,297 particles. For all 2D classification jobs, “align filament classes vertically” was turned ON. *Ab-inito* job with default settings resulted in a map that was used for initial refinement with or without symmetry applied. C6 refinement produced a high-resolution map that was used to identify initial helical parameters using a “symmetry search” job with a starting guess for rise of 30-50 and for twist of 1-30. The search returned a 41.8 Å estimation for rise and an 18° estimation for twist. A helical refinement job was started with a C6 initial volume and the estimated parameters, producing a final 3.22 Å map.

For masked refinements, we used masks generated in UCSF Chimera (v 1.13.1) using the molmap command as follows: molmap sel 20 (where “sel” is selected atoms and 20 is the resolution in Å). Masks were imported into CryoSPARC and used for 3D classification jobs.

### Model building

We used the web-interface for DeepTracer to generate an initial model, which was used to identify the homologous sequence using BLASTp. CollabFold v1.5.5 was used for AlphaFold model generation as well as to generate a flexible domain dimer model. For the dimer model, the flexible domain sequence [155-245] was input twice and submitted with default settings, resulting in five similar predicted models. Manual model editing was done in Coot 0.9.4. Model refinement was done in Phenix 1.21.2, first for the monomer alone, and then for the 6-member ring with NCS restraints.

Final C6 helical refinement map and core tail mode were deposited as EMD-48226 and PDB:9MFE. Composite full-length model of a single tail segment can be found in supplemental information.

## Supporting information

Supplemental Model 1

## Supplementary methods

### Purification of PP7 VLPs

PP7 VLPs were expressed in BL21(DE3) *E. coli* cells (New England Biolabs). Briefly, a single transformant BL21(DE3) colony was inoculated into a 250 mL baffled flask containing 50 mL of 2YT medium and a final concentration of 50 μg/mL streptomycin. The culture flask was grown to saturation overnight in a shaking incubator at 37 °C and 185 rpms. The following morning, a 2 L baffled Erlenmeyer flask containing 500 mL of 2YT medium and antibiotic was seeded with 5-10 mL of the saturated starter culture. The large expression culture was then grown at 37 °C in a shaking incubator to an OD_600_ between 0.8-1.0, at which point protein expression was induced with the addition of a final concentration of 1 mM IPTG. The cell culture was then incubated for 4 hours at 37 °C before harvesting the cells by centrifugation at 6000 rpms in a JA-16.250 rotor (Beckman Coulter) for 10 minutes. Cell pellets were stored at -80 °C prior to protein purification.

VLPs were purified by first resuspending the entire cell pellet from a 500 mL culture in 50 mL of potassium phosphate buffer (0.1 M, pH 7.0). Resuspended cells were lysed by sonication in an ice/water bath for a total of 10 minutes using 5 second pulses of 30-50 W with 5 seconds of rest between pulses. Clarified cell lysate was obtained by centrifuging the lysed cells at 14,000 rpms for 10 minutes at 4 °C in a JA-17 rotor (Beckman Coulter). VLPs were precipitated out of the clarified lysate by adding a final concentration of 0.265 g/mL ammonium sulfate and incubating the solution on an end-over-end mixer at 4 °C for 1 hour. Precipitated solids were collected by centrifugation, and were resuspended in 3-5 mL of fresh phosphate buffer after decanting the supernatant. Residual hydrophobic contaminants were removed by organic extraction of the resuspended VLPs with an equal volume of a 1:1 *n*-butanol/chloroform mixture. The aqueous and organic layers were subsequently separated by centrifugation and the upper aqueous layer was removed by aspiration. VLPs were further purified from the aqueous solution by passing them through 10-40% sucrose gradients (prepared in the same 0.1 M, pH 7.0 potassium phosphate buffer) for 4 °C for 4 hours at 28,000 rpms in a SW-32 rotor (Beckman Coulter). Lastly, VLPs were concentrated via ultracentrifugation at 68,000 rpms for 2 hours in a Type 70Ti rotor (Beckman Coulter) at 4 °C. The resulting supernatant was decanted, and the clear protein pellet was resuspended in fresh 0.1 M, pH 7.0 potassium phosphate buffer to a final concentration of 2 mg/mL, as determined using a Bradford Assay Kit (Pierce).

## Supplementary figures

**Figure S1.**
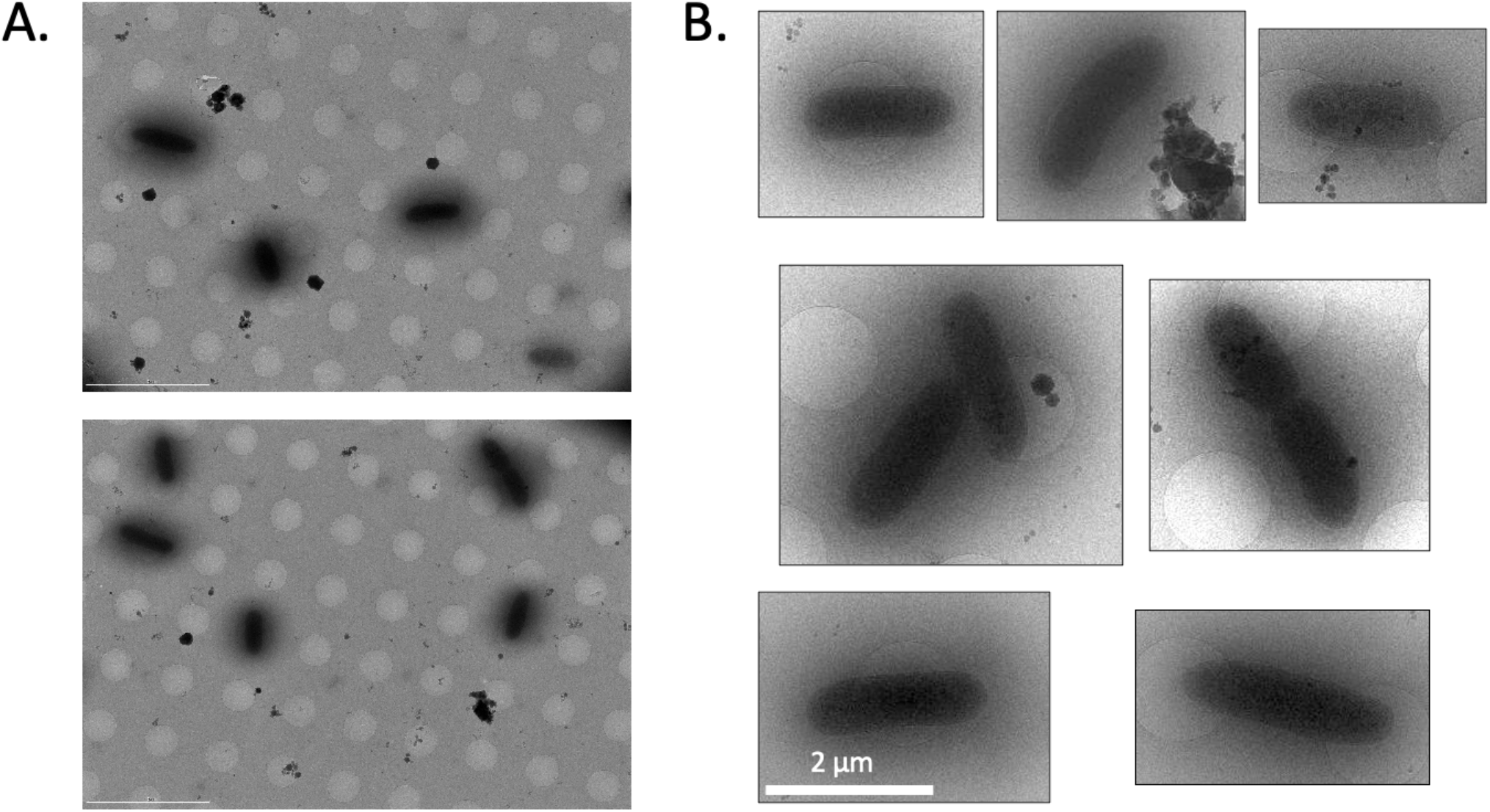
Bacterial contamination on the cryoEM plunge-frozen grid. A. Low-magnification images of the grid squares from the contaminated sample. Scale bar 5 μm. B. Cropped out bacteria from low-magnification images. Black and white levels were auto adjusted for to enhance detail visibility. Scale bar 2 μm.

**Figure S2.**
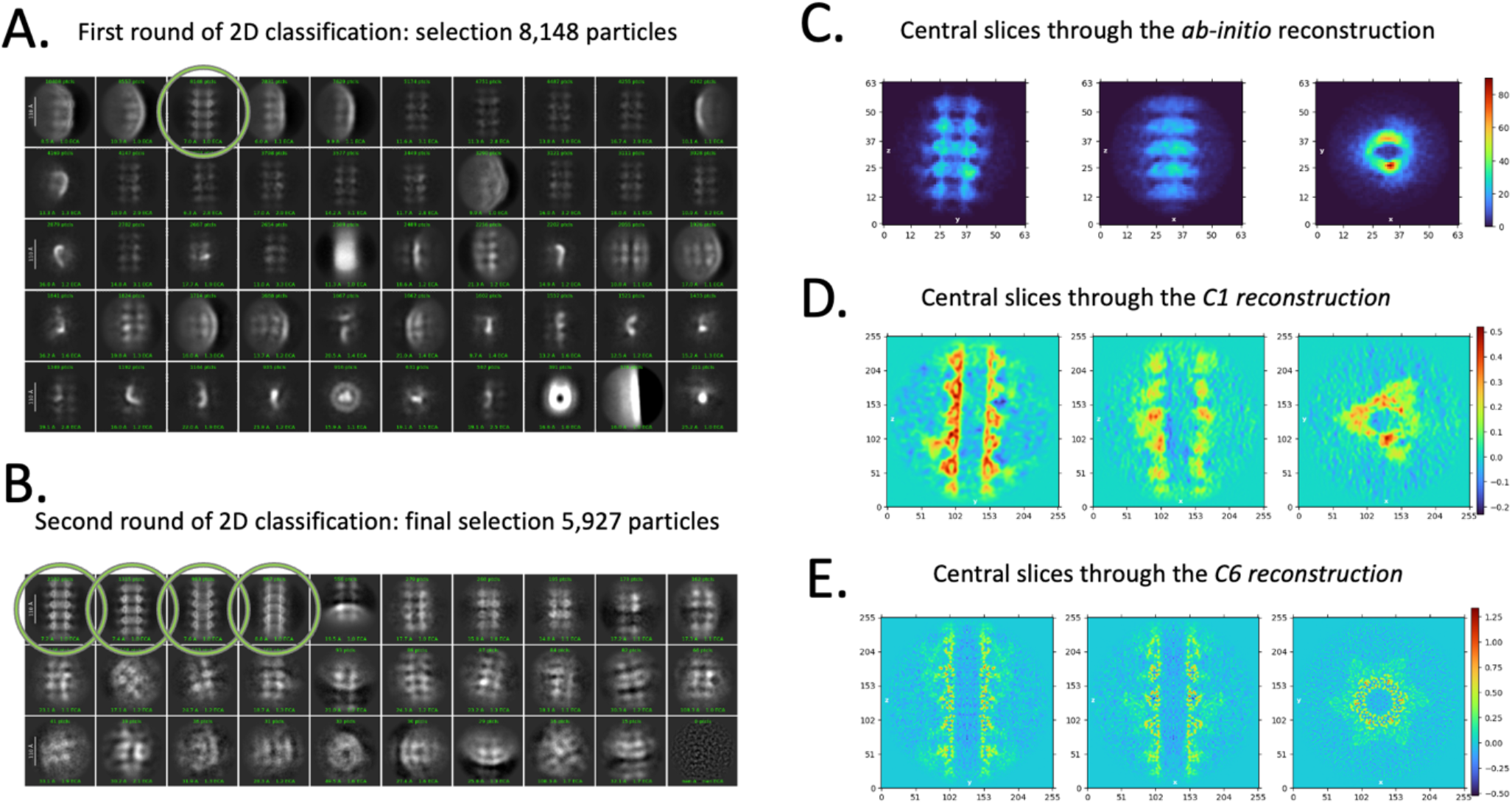
Data processing. A. Particles after template picking were extracted with the box size of 256 and 2D classified into 50 classes. B. Particles from the best-looking class in (A, green circle) with defined segments were selected for the second round of 2D classification, now into 30 classes. C. Particles from the four best-looking 2D classes (B, green circles) were selected to generate *ab-initio* reconstruction. D. C1 refinement using *ab-initio* model as input and particles from the second round of 2D classification. E. Same as D, but with C6 symmetry applied. Map from (E) was used to estimate helical parameters and was used as an initial volume for the final helical refinement (See Figure S3)

**Figure S3.**
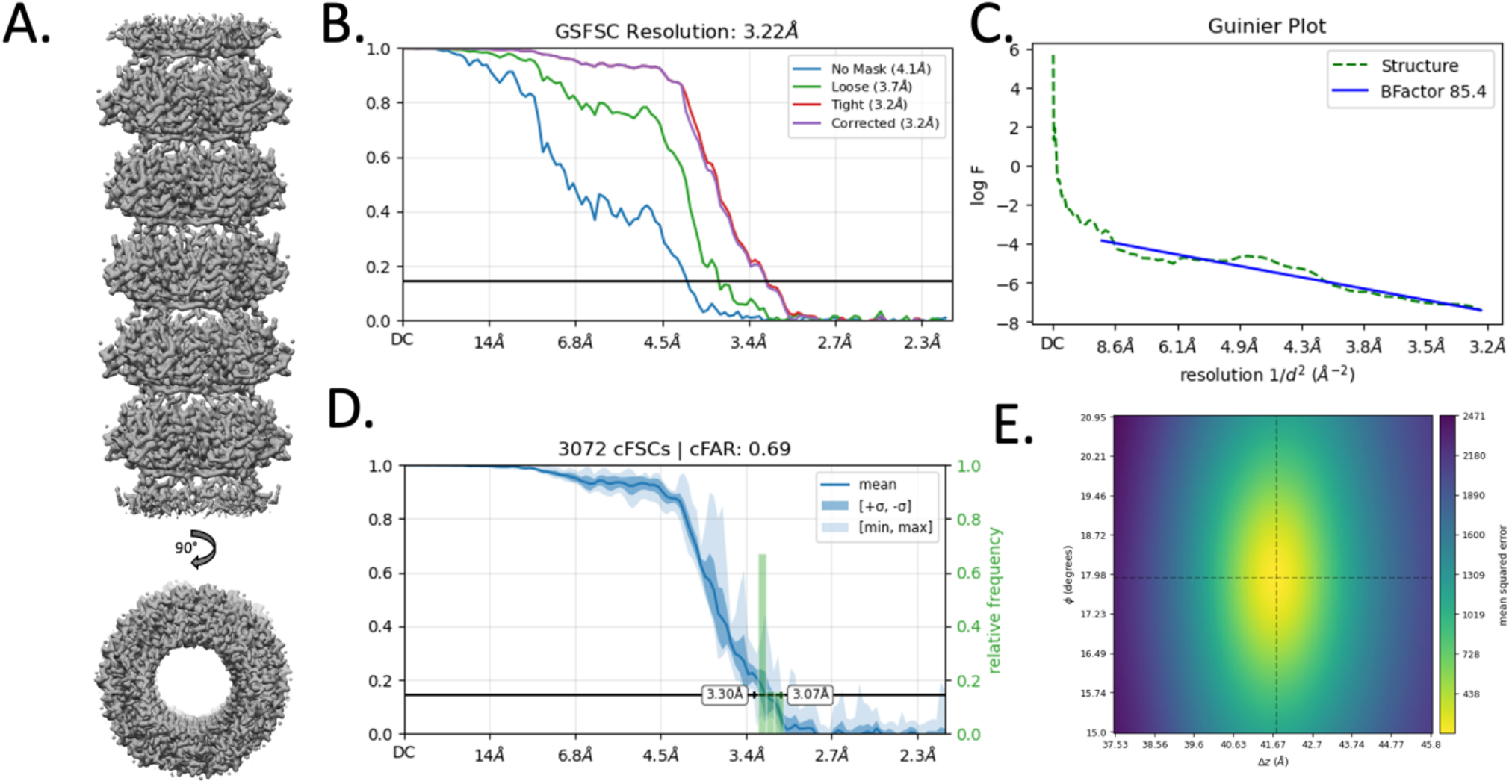
Final helical refinement details. A. Final map with helical and C6 symmetry applied. B. GSFSC plot showing the final map resolution of 3.22 Å. C. Guiner plot showing the b-factor estimation. B-factor correction of -85.4 was applied to the final map and the resulting map was used for structure prediction and model building D. Directional FSC plot showing isotropic resolution of the final map. E. Search plot for the final helical parameter estimation: twist 17.906, rise 41.765.

**Figure S4.**
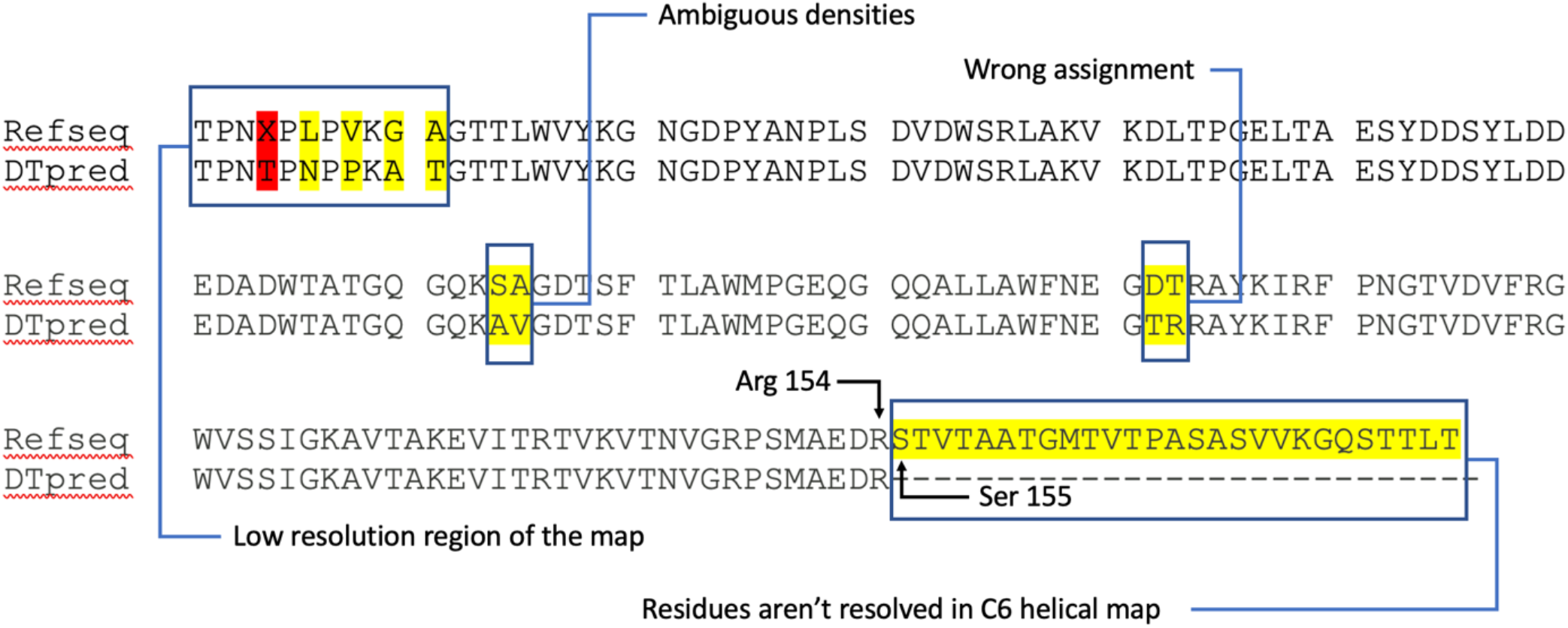
Comparison of the DeepTracer second run with the reference sequence. Highlighted in yellow and red are differences between the reference sequence and the one generated by the second run of DeepTracer. All mismatches were manually inspected in Coot and rebuilt to match the reference sequence. Locations of Arg154 and Ser155 are shown (numbering following the reference sequence).

**Figure S5.**
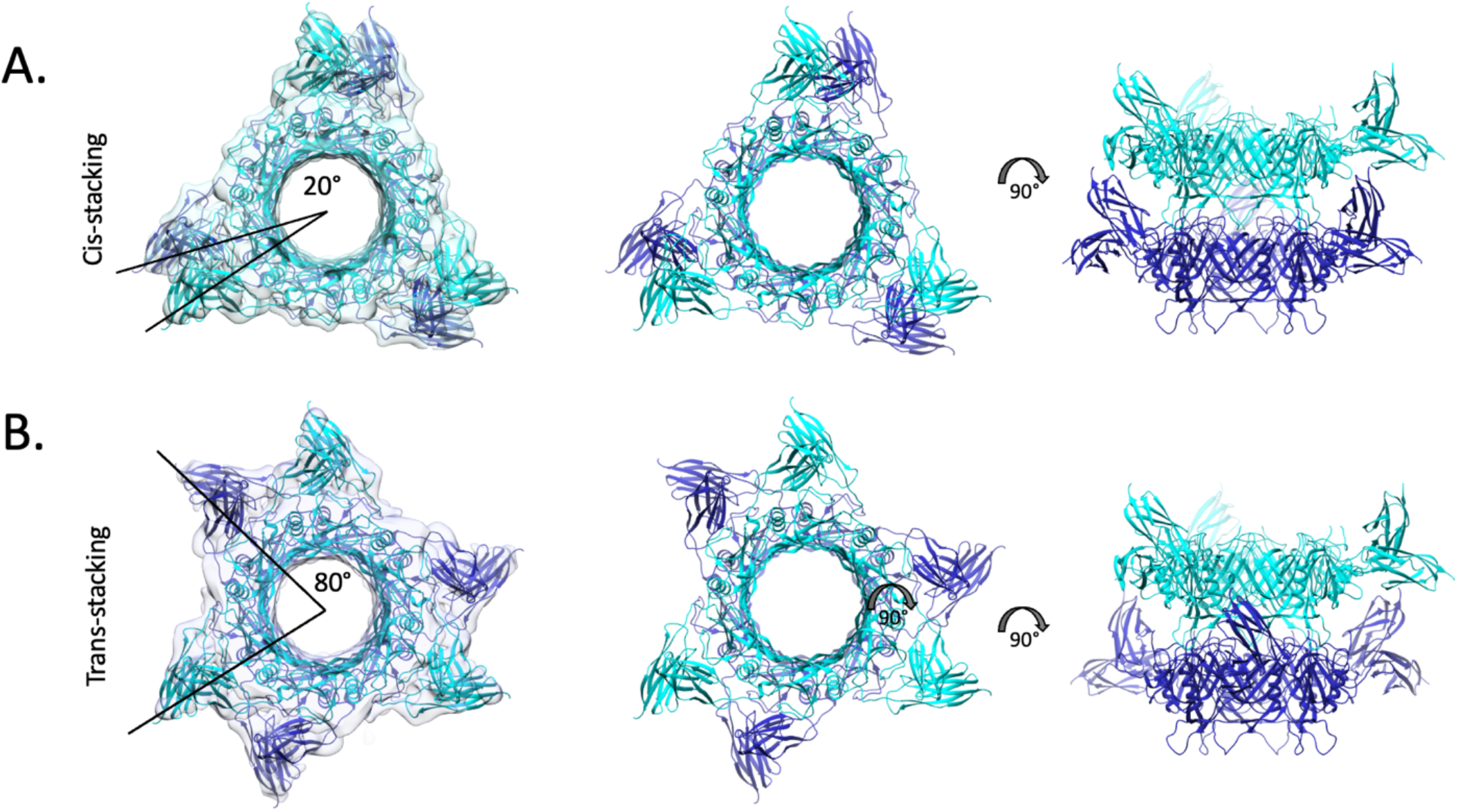
Two possible ways of segment stacking due to the symmetry mismatch. A. Cis-stacking, where the flexible domain dimers are positioned close to each other with ∼20 degrees offset. B. Trans-stacking, where the flexible domain dimers are positioned further away from each other with ∼80 degrees (or -40 degrees) offset.

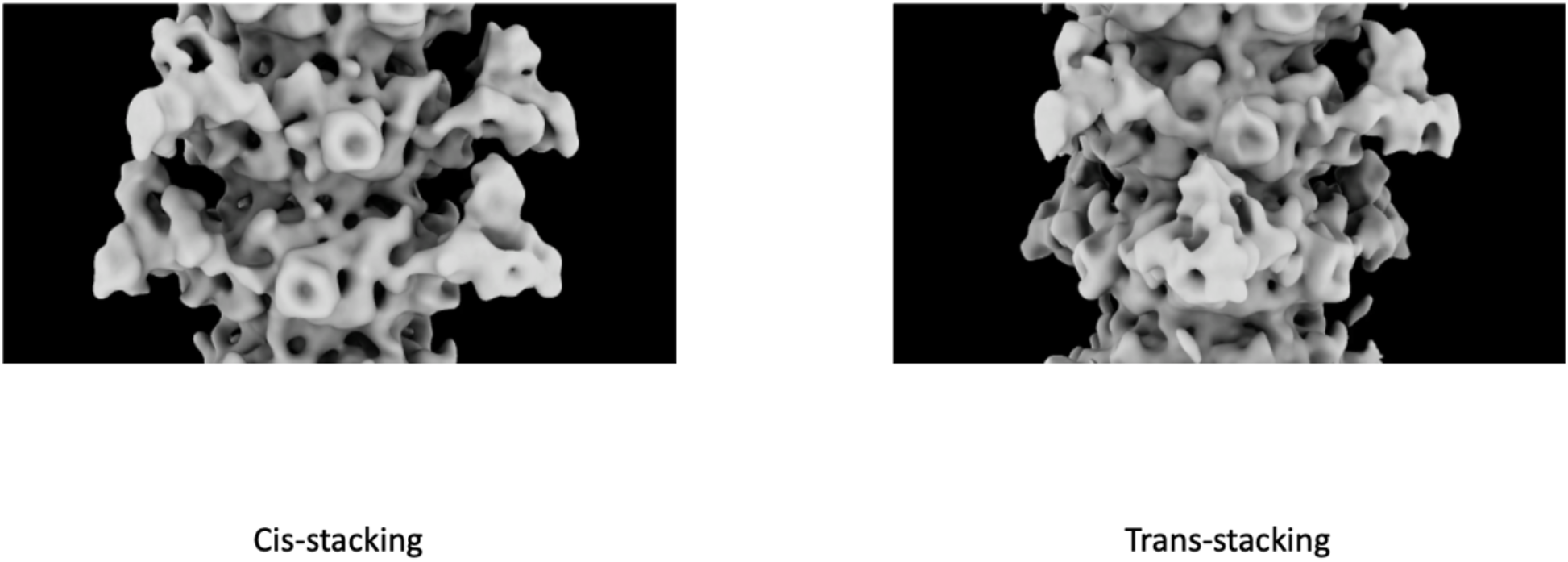

Supplementary videos are available for download.

Supplementary video 1. Cis-stacked segment.

Supplementary video 2. Trans-stacked segment.

